# Delineating the complex mechanistic interplay between autophagy and monocyte to macrophage differentiation: a functional perspective

**DOI:** 10.1101/2021.03.01.433396

**Authors:** Anindita Bhattacharya, Purnam Ghosh, Arpana Singh, Arnab Ghosh, Arghya Bhowmick, Deepak Kumar Sinha, Abhrajyoti Ghosh, Prosenjit Sen

## Abstract

Autophagy is an extremely essential cellular process aimed to clear redundant and damaged materials. In this study, we demonstrated that mTOR dependent classical autophagy is ubiquitously triggered in differentiating monocytes. Moreover, autophagy plays a decisive role in sustaining the process of monocyte to macrophage differentiation. We have delved deeper into understanding the underlying mechanistic complexities that trigger autophagy during differentiation. We have also shown that autophagy directs monocyte differentiation via protein degradation. Further, we delineated the complex cross-talk between autophagy and cell-cycle arrest in differentiating monocytes. This study also inspects the contribution of adhesion on various steps of autophagy and its ultimate impact on monocyte differentiation. Our study reveals new mechanistic insights into the process of autophagy associated with monocyte differentiation and would undoubtedly help to understand the intricacies of the process better for the effective design of therapeutics as autophagy and autophagy-related processes have enormous importance in human patho-physiology.

**Graphical Abstract:** 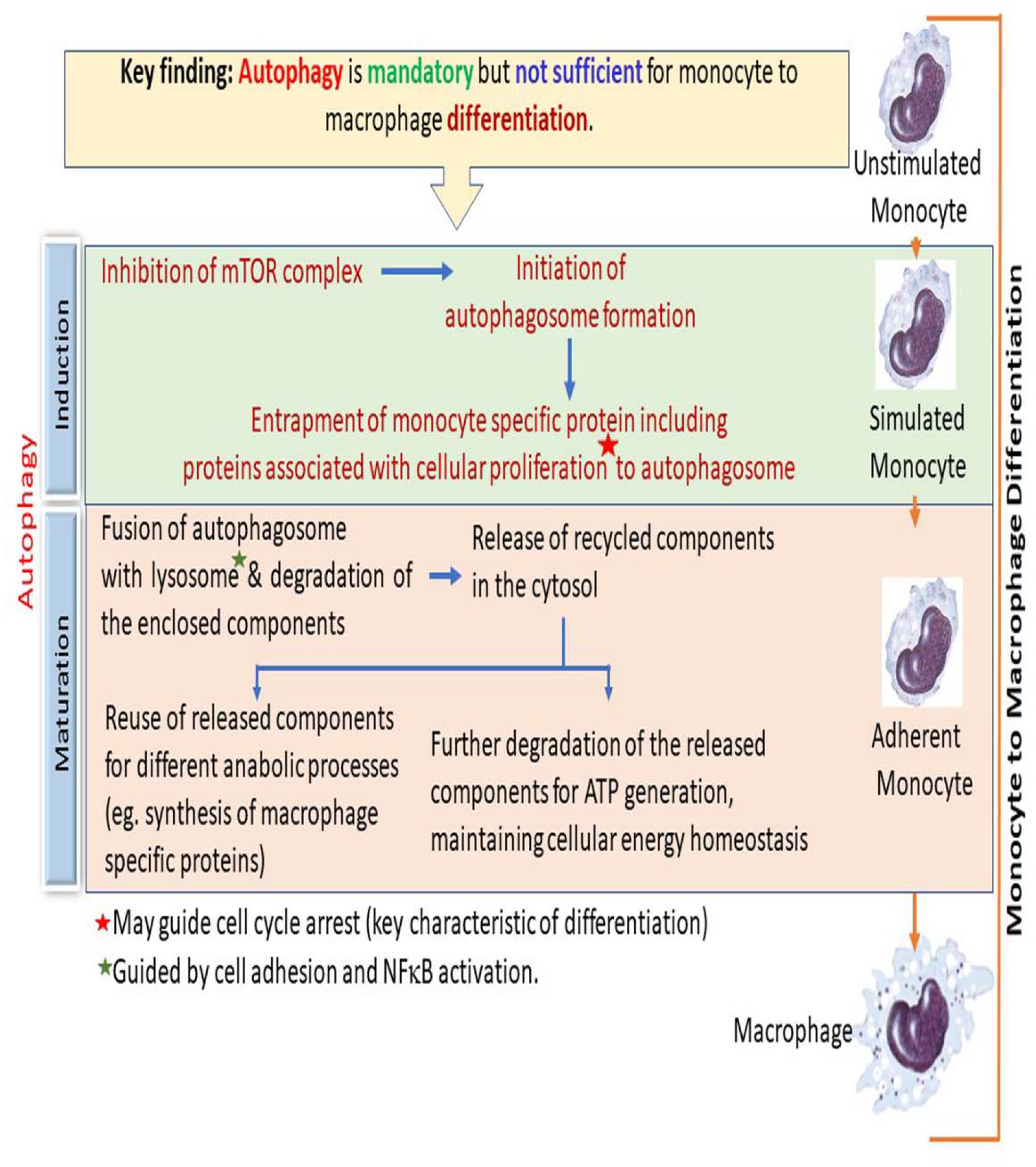

## 1. Introduction

Monocytes are bone-marrow-derived leukocytes that circulate in the blood and differentiate into macrophages by relocating from the bloodstream to the tissues. The mononuclear phagocyte system, a component of the innate immunity, is composed of blood-circulating monocytes and tissue-resident macrophages(Hume, 2006). During hematopoiesis in the bone-marrowhematopoietic stem cells differentiates to promonocytes. The promonocytes leave the bone-marrow, enter blood-circulation, and further differentiate into mature monocytes. In response to pro-inflammatory, metabolic, or immune stimuli, circulating monocytes infiltrate peripheral extravascular or inflamed tissues, where they differentiate into either macrophages or dendritic cells(Pittet et al., 2014).Yolk-sac derived tissue-resident macrophages replenish their pool by self-renewal, whereas blood-derived macrophages are maintained by a continuous supply of monocytes from the blood-stream(Gordon and Taylor, 2005),(Ginhoux and Jung, 2014),(Epelman et al., 2014). Macrophages occupy specific enclaves in organs, where they contribute to host-defense, development, tissue-remodeling and repair(Gordon and Plüddemann, 2017),(Das et al., 2015),(Rao et al., 2014).

Recent scientific developments strongly suggest that monocytes and monocyte-derived macrophages not only protect our system from invading pathogens but play a very important role in the development and prognosis of several diseases. The process of monocyte to macrophage differentiation is strictly regulated and any irregularity in this process may lead to life-threatening diseases like systemic sclerosis, Adult-Onset Still’s Disease (AOSD), primary biliary cholangitis, rheumatoid arthritis, etc (Giacomelli et al., 2018),(Bracaglia et al., 2017),(Kinne et al., 2000). Aberration during monocyte to macrophage differentiation may also cause several immune-deficiencies(Andrews and Sullivan, 2003),(Charles A Janeway et al., 2001),(Pathophysiology of Disease). Since the process of monocyte to macrophage differentiation has an immense significance on cellular physiology, it is important to unveil the underlying mechanistic details. Understanding the exhaustive immunological role of monocyte to macrophage differentiation will open new therapeutic windows.

Autophagy is a process involving the regulated and controlled mechanism of self–eating through which cells degrade unnecessary or dysfunctional components and reutilize the basic constituting macromolecules for future use(Glick et al., 2010). The process of autophagy starts with the formation of phagophores (Mizushima et al., 2002),(Hafner Česen et al., 2012) from the ER (endoplasmic reticulum), plasma membrane, mitochondria and ER mitochondria contact sites(Wei et al., 2018). This phagophore expands to engulf intra-cellular cargo of protein aggregates, organelles and ribosomes to forma utophagosome. Autophagosome matures through fusion with the lysosome(Mizushima et al., 2002) resulting in autophagolysosome, whereby promoting the degradation of autophagosomal contents by lysosomal acid proteases(Autophagy Gene List). Next, lysosomal permeases and transporters export the digested amino acids and other by-products of degradation back to the cytoplasm, where they can be re-used for different metabolic process. Thus, autophagy may be thought of as a cellular ‘recycling factory’ that also promotes energy efficiency through ATP generation and mediates damage control by removing non-functional proteins and organelles(Glick et al., 2010). Autophagy also plays a very important role in Xenophagy, ie the selective degradation of infectious foreign pathogens and hence, plays a very important role in innate immunity.

The ubiquitin-proteasome and autophagy-lysosomal pathways are the two major routes for protein and organelle clearance ineukaryotic cells. Ubiquitin-proteasome pathway predominantly degrades short-lived nuclear and cytosolic proteins(Lenk and Sommer, 2000),(Chowdhury and Enenkel, 2015),whereas the bulk degradation ofcytosolic proteins or organelles occurs through autophagy.

The classical pathway of autophagy is regulated by the serine/threonine protein kinase mTOR (Sridharan et al., 2011),(Jung et al., 2010) which negatively regulates autophagy. Normally in the condition of starvation, mTOR gets inhibited and autophagy gets initiated to recycle intracellular constituents and to provide an alternative efficient source of energy(Gallagher et al., 2016).Apart from this mTOR independent pathways are also there, namely, the inositol signaling pathway(Sarkar, 2013), Ca2^+^/Calpain pathway (Gordon et al., 1993) cAMP/Epac/Ins(1,4,5)*P*3 pathway(Noda and Ohsumi, 1998).

In addition to the metabolic role, autophagy also regulates the immune system of our body. Along with phagocytosis, autophagy also has a crucial role in the engulfment and disposal of unwanted, and potentially harmfulmaterials(Germic et al., 2019).Autophagycan also promote mycobacterium clearance by stimulating phagosomal maturation (Dengiel et al., 2005),(Armstrong and Hart, 1975)and through the production of antimicrobial peptides(Alonso et al., 2007; Ponpuak et al., 2010). Autophagy possesses a protective role against *Toxoplasma gondii* infection(Martens et al., 2005; Zhao et al., 2008). Along with having an antiviral property against herpessimplex virus type 1 (HSV-1) and Sinbis virus(SIN)(Kuballa et al., 2012) autophagy is also important in controlling the spreadof fungal infection and susceptibility to disease in plants(Dengiel et al., 2005). It also facilitates antigen presentation via MHC class II molecules to CD4+ T cells(Dengiel et al., 2005) and modulates neutrophil activity by regulating antibody-dependent cellular cytotoxicity (ADCC) of cancer cells andthe formation of neutrophil extracellular traps (NETs) bywhich neutrophils capture microbes(Kuballa et al., 2012). NETs are antimicrobial extracellular chromatinstructures released by neutrophils undergoinga form of cell death termed NETosis (Brinkmann et al., 2004; Fuchs et al., 2007). Autophagy is also reported to be associated with B/T lymphocytes, dendritic cells, macrophages and NKT cells survival and development(Kuballa et al., 2012).

Moreover, autophagy is also reported to be associated with several kinds of cellular differentiation processes including that of erythrocytes, lymphocytes, and adipocytes.(Miller et al., 2008; Pua et al., 2009; Sandoval et al., 2008; Singh et al., 2009; Stephenson et al., 2009; Zhang et al., 2009). There are few literature reports correlating autophagy with monocyte to macrophage differentiation process.Circulating monocytes have a very short life span and they are destined to undergo apoptotic cell death if failed to differentiate(Furth and Colin, 1968),(Whitelaw, 1972),(Ricardo et al., 2008).Cytokines like GM-CSF andM-CSF (Ricardo et al., 2008)as well as inflammatory stimuli such as Lipopolysaccharide (LPS), Phorbol-12-myristate-13-acetate (PMA) (Bhattacharya et al., 2020)and 1,25-dihydroxyvitamin D3(VD3)(Arboleda Alzate et al., 2017), can induce their differentiation into macrophages(Ricardo et al., 2008). However,detailed molecular events have not been elucidated to date to provide mechanistic insight into the differentiation process. Recently, Yuk et al. have shown thatVD3, induces autophagy in monocytes(Yuk et al., 2009). Jacquel et al have also found that autophagy is required for CSF-1–induced macrophagic differentiation and acquisition of phagocytic functions(Jacquel et al., 2016). Zhang et al. have reported that autophagy induction is essential for GM-CSF triggered monocyte-macrophage differentiation. They have also established that autophagy is a prerequisite for protecting the cells from apoptosis(Zhang et al., 2012). However, the mechanism and the purpose of induction of autophagy during monocyte-macrophage differentiation is not well elucidated.

Through this study, we have tried to explore the complicated crosstalk between autophagy and monocyte differentiation. Apart from establishing the significance of mTOR dependent autophagy as a crucial regulator of monocyte differentiation, we have also instituted the detailed correlationbetween cellular adhesion and autophagy in the context of monocyte to macrophage differentiation. We have delved deeper into understanding the functional parameters of autophagy involved in the process of monocyte differentiation and the probable mechanistic interpretation portrayed in this study will undoubtedly provide a platform to visualize the differentiation process from the molecular level. Moreover, the autophagic pathways can be targeted to develop new therapeutics against differentiation-related immune disorders.

## 2. Materials & Methods

### 2.1 Cell culture

Monocytic leukemia cell line (THP-1) of ATCC origin was acquired from Dr. Amitabho Sengupta, IICB as a kind gift. THP-1 was cultured in RPMI-1640 medium supplemented with 10% heat-inactivated FBS and 5% Penstrep (Invitrogen and Himedia) in an incubator fixed at 37°C and equipped with 5% CO_2_ supply. For triggering monocyte to macrophage differentiation PMA (50ng/ml) or LPS (100μg/ml) were used and stimulated THP-1 cells are seeded either on a confocal dish (for microscopy) or on a commercially available adhesion compatible dish and kept in the incubator for requisite time point. Human peripheral blood mononuclear cells (PBMC) were isolated by density gradient centrifugation protocol using ficol (Himedia) gradient. Blood samples from healthy human volunteers were collected in (10% V/V) citrate buffer. The freshly collected blood was half diluted with PBS and layered over the top of ficol. Next, the sample was centrifuged at 400g for 20 minutes to obtain the buffy coat containing PBMC. The buffy coat was washed twice with PBS and subsequently dissolved in RPMI-1640, seeded into a Petri dish that was incubated overnight. The next day, the suspended neutrophils and lymphocytes were washed with supplementation of fresh medium to the adhered PBMC.

### 2.2 Western blot

Western blot was performed following the standard protocol. Cell-lysis was done with Laemmeli buffer. Following that lysates were loaded and separated in SDS PAGE. Next, the electrophoresed proteins were transferred from gel to PVDF membranes (Millipore) following with incubation with primary and secondary antibodies and finally bands were detected. Pre-stained molecular markers (Biorad) were used to estimate the molecular weight of samples. The protein expression level was quantified with Image J software (NIH).

### 2.3 RT-PCR

Trizol (Life-technologies) was used for total RNA isolation. Reverse transcription was done from the extracted mRNA with life technologies kit. Next, desired gene was PCR amplified using that cDNA as template for quantitative estimation of target gene transcription. β-actin was used as an internal control.

### 2.4 Phagocytosis assay

200nm (Invitrogen) carboxylated latex fluorescent beads were added to the cells and incubated for two hours. At the end of incubation, the cells were washed with PBS to remove the non-phagocytosed beads. RFP expressing *E.coli* culture was added to the cells. 30 minutes later the cells were washed with PBS to remove the free bacteria. Fluorescence Z-stack images of the cells were acquired. Maximum intensity projection image from the previously acquired Z-stacks was obtained to count the phagocytic cell percentage. The fluorescence intensity of z-projected images was scaled such that the background intensity becomes 1. The cells with average fluorescence intensity more than 10% of the background were considered as phagocytic.

### 2.5 NO measurement

Griess reagent was incubated with spent medium and absorbance was measured at 540 nm.

### 2.6 ROS measurement

25μM DCFDA (Sigma) was incubated with cells for 30 minutes. Next, the cell-lysis was done with DMSO and the fluorescence was measured at Ex485 nm/Em535 nm.

### 2.7 Immunostaining

Cells were washed with PBS and fixed with 4% paraformaldehyde in PBS for 10 min. Next the cells were permeabilized with 0.1% Triton x 100 for 2 mins (in case of LC3 staining), washed with PBS and blocked with 5% BSA for an hour. Primary antibody (Abcam/CST) incubation was done overnight, washed with PBS and then incubated with Alexa tagged secondary antibody (CST) for 45 minutes, finally washed with PBS, counterstained with Hoechst and imaged. Cells were imaged with the sCMOS camera (Orca Flash 4.0, Hamamatsu) on an inverted fluorescence microscope from Carl Zeiss (Axio-observer Z1) and a confocal microscope (Carl Zeiss LSM880). Image J was used for image processing.

### 2.8 CYTO-ID staining

1μl of Cyto-ID Green Detection Reagent (Enzo-lifesciences kit) was incubated with 10^5^ to 10^6^ cells/ml for 30 min under standard tissue culture conditions at 37 °C, 5% CO_2_ in the dark. Then the cells were washed with PBS and imaged at 488nm excitation filter.

### 2.9 DQ™-Red BSA staining

DQ™-Red BSA (10 μg/ml) was added to cells following incubation for 4 hours at 37 °C. Finally the cells were washed with PBS to remove the unbound stain and imaged at 590 nm excitation filter and 620 nm emission filter.

### 2.10 Quantification of total cellular protein

Cells were lysed using RIPA buffer (50mM TRIS, pH 8, 150mM NaCl, 1% NP-40, 0.5% Sodium deoxycholate, 0.1% SDS). The lysate was incubated with Bradford reagent for 20minutes at 37° C. Absorbance was measured at 595nm.

### 2.11 Estimation of cellular ATP level

Cellular ATP content was assessed using Promega ENLITENATP kit. Cells were washed with ice-chilled PBS and lysed with ice-cold Phenol TE by homogenization in ice. The cell lysate was mixed with chilled chloroform and water followed with shaking and centrifugation at 10000g for 5 min at 4°c. The supernatant was diluted with water and mixed with rL/L reagent of kit. The ATP content was read at 560 nm using a luminometer (Thermo Scientific Luminoskan Ascent)

### 2.12 Flow cytometry

Primarily, cellular enumeration has been performed using flow cytometry based analysis. For cell cycle analysis standard protocol has been followed. Following harvesting, the cells were washed with ice-cold PBS twice. Next, cells were fixed with 70% ethanol while keeping the cells on ice. After 1 hour, the cell suspension was centrifuged, the supernatant was discarded. Pellets were washed twice with PBS. Next, the cell pellet was dissolved in PBS followed with 30mins incubation with RNAse A (100μg/ml). Finally, Propidium Iodide (PI) (5 μg/ml) was added and kept for 15 minutes and subjected to FACS analysis (BD FACSAria).

### 2.13 Cell labeling with biotin and immunoprecipitation

This experiment was carried out following the protocol described by Tarradas et al (Tarradas et al., 2013). Keeping the cells on ice growth medium was removed. Cells were washed with ice-cold PBS and incubated for 30 min on ice in the cold room with gentle rocking with biotin solution (2.5 mg/ml biotin reagent in DPBS). Cells were washed thrice for 5 minutes by 1 ml of cold 100 mM Glycine in DPBS, on icein the cold room with gentle rocking to neutralize the unbound excess biotin. Next, One set of cells were treated with PMA (50ng/ml) and incubated for differentiation. Other set of cells were instantaneously lysed. Cells were lysed with lysis buffer (50 mMTris/HCl (pH 7.4), 150 mM NaCl, 1 mM EDTA, 1% (w/v) Triton X-100& protease inhibitor). For each sample 40 μl of 50% slurry of Immobilized/ NeutrAvidin Ultralink beads were taken, washed twic ewith PBS and twice again with wash buffer (50 mMTris/HCl (pH 7.4), 150 mM NaCl, 1 mM EDTA, 1% (w/v) Triton X-100) After the last wash, the precipitated beads were resuspended in lysis buffer. Lysates were spun at 16,000 x g (maximum speed) for 15 min at 4 °C.A portion of supernatants was stored at −80 °C for input. The remaining supernatant was incubated with the NeutrAvidin beads for overnight in the rotating wheel at slow speed in the cold room. The next day, samples were centrifuged at 16,000 x g, 30 sec at 4 °C. Beads were washed with lysis buffer and wash buffer. Pellets were resuspended in of 2x laemelli buffer and heated at 95 °C for 10 min. Samples were analyzed by SDS-PAGE and Western blot

### 2.14 Isolation of autophagosome

Autophagosome purification was modified from the protocol described in Ref. 111(Chen et al., 2015)Cells were washed in PBS and lysed by homogenization. The post-nuclear supernatant was further centrifuged at 10,000gor 20 min. The supernatant was centrifuged at 10,0000*g* for 90 min. The size of isolated autophagosome was confirmed by DLS study and LC3II enrichment also suggested successful purification of autophagosome fraction.

### 2.15 Cell starvation in minimal buffer

Cells were starved in Minimal Buffer Media EBSS (CaCl_2_ 0.2 g/L, KCl 0.4 g/L, MgSO_4_ 0.097 g/L, NaCl 6.80 g/L,NaHCO_3_ 2.2 g/L, NaH_2_PO_4_ 0.14 g/L,and Glucose1g/L) and HBSS (NaCl 8 g/L, KCl 0.4 g/L,CaCl_2_ 0.14 g/L, MgSO_4_ 0.1 g/L, MgCl_2_ 0.1 g/L, Na2HPO_4_ 0.06 g/L, KH_2_PO_4_ 0.6 g/L, NaHCO_3_0.35 g/Land Glucose 1 g/L).

### 2.16 Proteomics study

For proteomic analysis, a combined approach has been adopted by using MALDI-MS and LCMS-MS.

For broad-scale detection of the target protein, MALDI-TOF-MS was used. Here, the pellet was washed with 70% ethanol followed by centrifugation at 16000 g for 10mins. After removal of the supernatant, the pellet was dried and solubilized in 20μl of 10% formic acid and mixed with an equal volume of saturated sinapinic acid solution (40mg in 1ml of 60%acetonitrile/37% H_2_O/3%TFA). Next, spotting was done for each sample in triplicate on a MALDI TOF plate and reading was taken. Finally, from the obtained datasheet of m/z ratio, the peaks (with optimum intensity) were tallied with the masses of expected proteins.

Next, for LC-MS-MS based analysis, the pellet was dissolved in 25mM ammonium bicarbonate buffer followed by probe sonication for lysis. Then protein concentration was quantified in a different sample using BCA/Bradford methods of protein estimation. Next, approximately 200μg of protein was taken from each of the samples and mixed with 100 μl of 10mM DTT (made in 25mM ammonium bicarbonate buffer). Incubation was carried out at 56°C with shaking at 300rpm for 1hour. All the samples were cooled at room temperature for 15 mins. Next, 200 μl of 55mM iodoacetamide (made in 25mM ammonium bicarbonate buffer) was added to it followed by incubation at 25°C with shaking at 300rpm for 45 mins in dark. For precipitation of the proteins, 600 μlof 50% (W/V)TCA was added to each sample and incubated on ice for 2hour. Next, samples were centrifuged at 13000rpm for 15mins at 4°C. After removal of the supernatant, pellets were washed with 1ml cold 25mM ammonium bicarbonate buffer, vortexed briefly, and centrifuged at 13000rpm for 5mins at 4°C. After removal of the supernatant, pellets were dissolved in 100 μl of 50mM ammonium bicarbonate buffer. Next, 5 μl of sequencing grade trypsin was added to each sample and overnight incubation was carried out at 37°C with mild shaking. Next, the digested samples were dried using speedvac and the pellets were redissolved in 0.1%FA/3%ACN. Finally, the sample was injected into the LC-ESI-MS column. After detection, the data was analyzed using Progenesis QI software for proteomic analysis.

### 2.17 Animal Study

Indian Association for the Cultivation of Science (IACS) Institutional Animal Ethics Committee approved all the mice related studies. 3MA and Chloroquine were introduced into 6-wk-old male BALB/c mice (NIN; Hyderabad, India) via tail-vein injection (7 mice were taken in each group). 48 hours post-injection, 10^8 RFP-expressing *Escherichia coli* cells were injected into the peritoneal cavity of these mice and were kept for another 48 hours. Euthanization was performed by rapid cervical dislocation. Next, 5ml of chilled RPMI-1640 was injected in both sides of the peritoneum with consequent aspiration of the peritoneal fluid. Peritoneal exudate cells (PEC) were spun at a refrigerated centrifuge for 10 min at 400 × *g*. The supernatant was discarded and the pellet was dissolved in RPMI-1640 medium by gently tapping the bottom of the tube and pipetting up and down. Finally, an equal volume of cell suspension was seeded for each treatment. The cells were allowed to adhere to the substrate by centrifuging them for 1 to 2 hr at 37°C. Nonadherent cells are removed by gentle washing three times with warm PBS. 2 days post-seeding cell enumeration was performed by staining the cells with Hoechst (Sigma)(110ng/ml). Following staining, the cells were lysed with DMSO. Fluorescence intensity was measured at 361nm/497nm. For estimation of bacterial load in peritoneal exudate, it was treated with 0.5% sodium deoxycholate, followed by spreading on Ampicillin containing plate. The rest of the experiments were carried out according to the standard protocols described previously. Finally, the sizes of the spleen were estimated (measurement has been performed in three dimensions, a, b and c). All studies on the animal model were performed abiding the Declaration of Helsinki principles.

### 2.18 Statistical analysis

All the techniques applied here are representative of at least three (n≥3) independent experiments. The data presented here are as mean ±S.E of the mean and the differences are considered to be statistically significant at p < 0.05 using the student’s t-test. Graphpad prism was used for statistical analysis.

## 3. Results

### 3.1 Autophagy is induced during monocyte to macrophage differentiation

We examined whether autophagy is induced in monocytes in response to chemical stimuli of differentiation, namely LPS and PMA.THP-1 cells were treated with LPS and PMA to induce their differentiation to macrophages.Conversion of cytosolic diffused LC3 to processed and punctate LC3 characteristically represents the initiation of an autophagic response(Barth et al., 2010). In unstimulated monocytes, a diffused pattern of LC3 was found whereas, a significant increase in the processed LC3-IIB was observed in LPS/PMA stimulated cells (Fig. 1A and B). This observation was further validated by CYTO-ID staining (Fig. 1C)(Picot et al., 2019).Next, we assayed autophagy maturation (fusion of the autophagosome with lysosome) in differentiating monocytes by DQ Red BSA staining(Zhang et al., 2012). The fluorescent puncta of DQ Red BSA started appearing 6-8 hours after incubating the cells with LPS/PMA; reinforcing our claim that autophagy is induced in differentiating monocytes (Fig. 1D). Previous work from our group had established that adhesion is necessary and sufficient to drive monocyte differentiation in the 2D/3D microenvironment in the absence of any chemical stimuli(Bhattacharya et al., 2018). Therefore,the promotion of adhesion can drive spontaneous differentiation. In this scenario, we examined whether autophagy induction during monocyte differentiation was a specialized phenomenon occurring only in the presence of any chemical stimuli or, a much more generalized event associated with differentiation. THP-1 cells were seeded on collagen and fibronectin-coated glass surface and incubated for the requisite period. Adherent cells were found to contain processed LC3 (Fig. 1B, Sup Fig. 1A). Our data confirm autophagy induction and maturation in spontaneously differentiating monocytes seeded on collagen and fibronectin (Fig. 1B, Sup Fig. 1A-C).We further validated these observations in human blood-derived monocytes (PBMCs). Interestingly, we found similar induction and maturation of autophagy in differentiating PBMCs irrespective of the mode of induction (Sup Fig. 1D-G). Overall, our data suggest thatautophagy is ubiquitouslyinduced during monocyte to macrophage differentiation independent of the external stimuli.

**Fig. 1.**
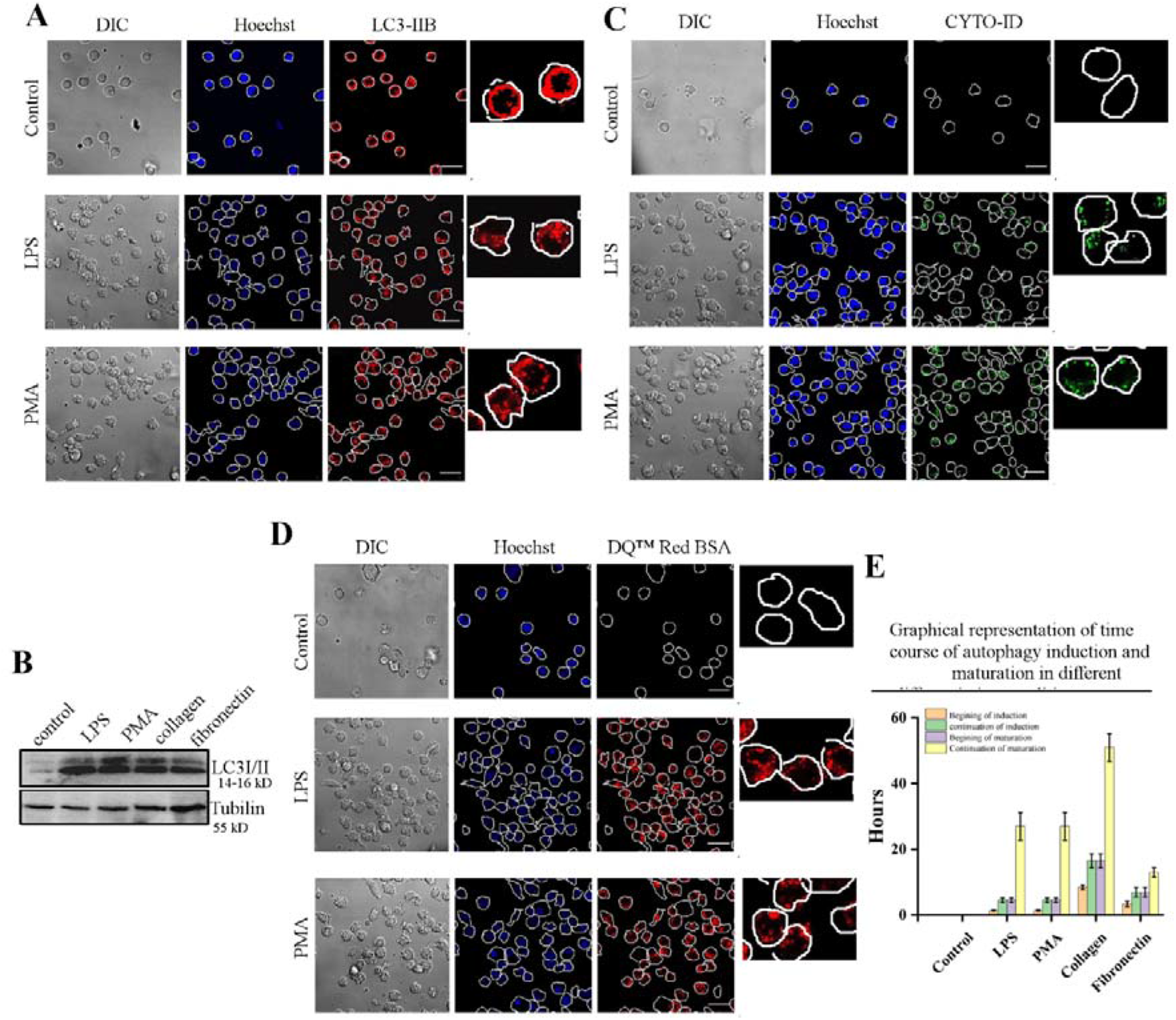
Autophagy is induced during monocyte to macrophage differentiation. (A) DIC (left panel) and fluorescent images of Hoechst stained (middle panel) and LC3-IIB (right panel) immunostained control (upper), LPS (middle) and PMA (lower) stimulated THP-1 cells. Magnified view of LC3-IIB immunostained cells are represented at the right side of the panel.(B) Western Blot of LC3 I/II in control; LPS, PMA treated and Collagen and Fibronectin surface-seeded THP-1 cells. Tubulin is used as the loading control. (C) DIC (left panel) and fluorescent images of Hoechst (middle panel) and CYTO-ID (right panel) stained control (upper), LPS (middle) and PMA (lower) stimulated THP-1 cells. Magnified view of CYTO-ID stained cells are represented at right side of the panel. (D) DIC (left panel) and fluorescent images of Hoechst (middle panel) and DQ™ Red BSA (right panel) stained control (upper), LPS (middle) and PMA (lower) stimulated THP-1cells. Magnified view of DQ™ Red BSA stained cells are represented at right side of the panel. (E) Graphical representation of time-course of autophagy induction and maturation under different conditions of differentiation viz. LPS, PMA, Collagen and fibronection. Scale Bar 20 μm. Graphical analyses are performed with standard error bars after repeating the experiment at least thrice.

### 3.2 Autophagy is induced via the mTOR pathway in differentiating monocytes

Next, we aimed to investigate the detailed mechanism involvedinthe process of autophagy accompanyingmonocyte differentiation. Mechanistic target of rapamycin (mTOR) is a key player of nutrient sensing and signaling(Ballou and Lin, 2008). There is a dynamic signaling reciprocity between mTOR and autophagy; and mTOR functions as the principal homoeostatic governor of cell growth, regulating whether anabolic or catabolic reactions are to be favored(Ballou and Lin, 2008; Saxton and Sabatini, 2017),(Dunlop and Tee, 2014).To check the involvement of mTOR dependent autophagic pathway during monocyte differentiation, we assayed the phosphorylation status of mTOR in monocytes stimulated with LPS/PMA. We observed significantly diminished phosphorylation of mTOR, indicative of the involvement of an mTOR dependent autophagy pathway during monocyte differentiation (Fig. 2A).Further, to examine whether constitutive activation of mTOR,i.e. inhibition of classical autophagy can affect the process of monocyte differentiation, we treated the monocytes with Phosphatidic acid (PA) before stimulation. PA is known to stabilizemTOR by binding to the FRB domain, competing with the inhibitory proteins and stabilizing as well as allosterically activating mTORC1(Fang et al., 2001; Yoon et al., 2011).Interestingly, we observed phosphatidic acid treatment seriously hampered monocyte differentiation (Fig. 2B-D). Similar mTOR mediated autophagy is also involved in differentiating PBMCs as indicated by the PA mediated inhibition of PBMC differentiation(Sup Fig. 2).Overall, our data suggest that classical mTOR dependent autophagy is induced during monocyte to macrophage differentiation, acting as a crucial regulator of the differentiation process.

**Fig. 2.**
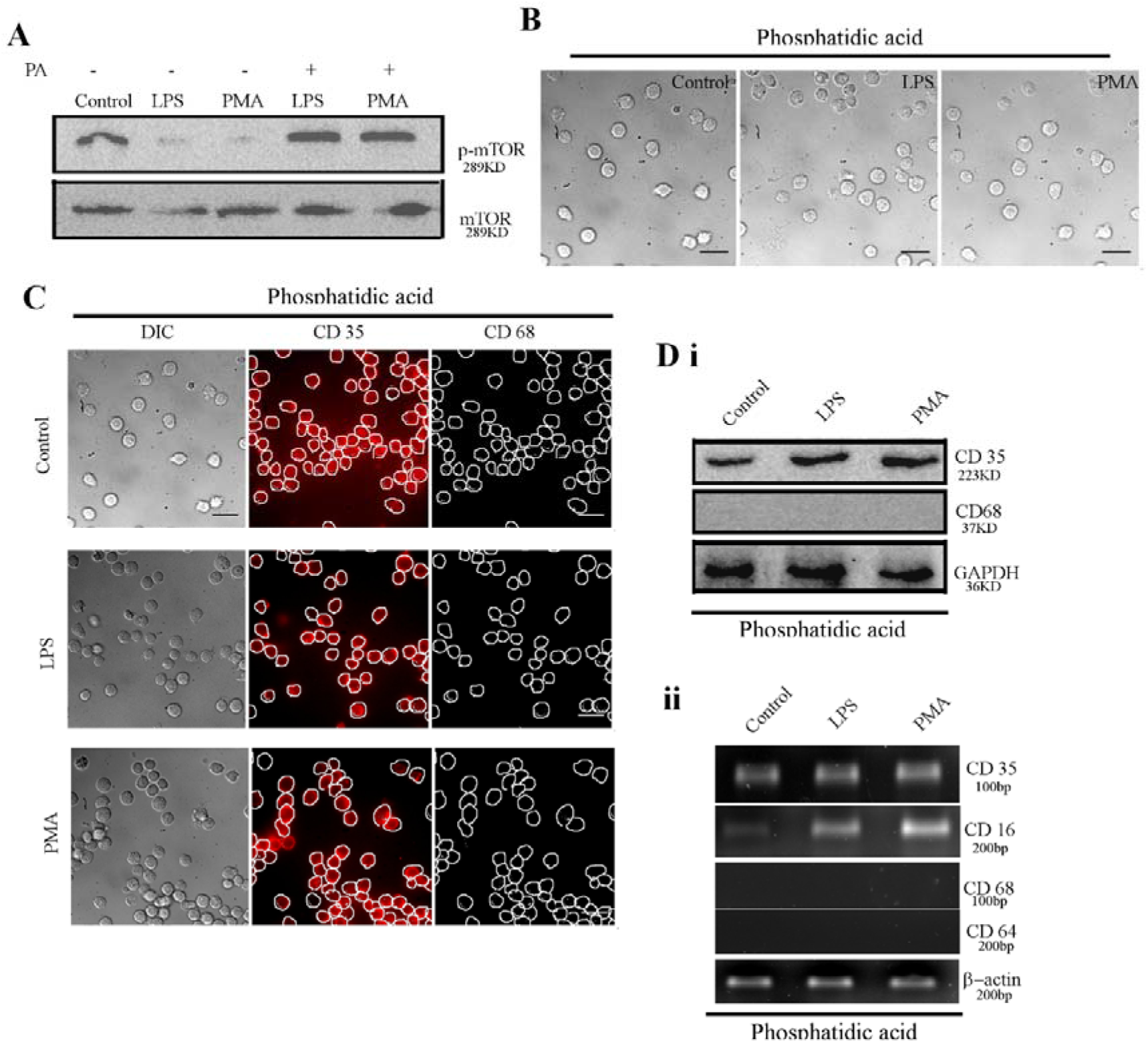
Autophagy is induced via mTOR pathway in differentiating monocytes. (A) Western Blot of phospho and non-phospho m-TOR of control; LPS, PMA stimulated THP-1 cells in absence (lanes 1-3 from left) and presence (lanes 4-5 from left) of Phosphatidic acid (PA). (B) DIC images of control (left), LPS (middle), PMA (right) stimulated THP-1 cells in presence of Phosphatidic acid. (C) DIC (left panel) and immuno-fluorescence images of CD35 (middle panel) and CD68 (right panel) markers of Phosphatidic acid treated control (upper), LPS (middle) and PMA (lower) stimulated THP-1 cells. (D) Western Blot (i) and Semi-quantitative RT-PCR analysis (ii) of monocyte and macrophage markers of control; LPS, PMA stimulated THP-1 cells in presence of Phosphatidic acid. GAPDH (in western blot) actin (in RT PCR) are used as a loading control. Scale Bar 25 μm.

### 3.3 Autophagy promotes monocyte to macrophage differentiation

Consequently, we wanted to investigate whether autophagy and monocyte differentiation are two intimately co-related or unrelated parallel pathways.Todetermine the effect of autophagy on differentiation, we pretreated the monocytes with different autophagy inhibitors; namely 3-Methyl Adenine (3-MA), Chloroquine (CQ), adenoviral construct of Rab7 Dominant Negative mutant (Rab7 DN) and Bafilomycin followed by chemical stimulation to block the process of autophagy at different stages. On studying the morphology, we observed that 3-MA treated cells remained in the suspension state, whereas CQ, Rab7DN and Bafilomycin treated cells adhered loosely, remained spherical and did not assume the shape characteristic of macrophages (Fig. 3A,Sup Fig. 3A). Subsequent marker analysis revealed that 3-MA treated cells expressed monocyte markers but cells treated with CQ, Rab7 DN and Bafilomycin expressed the macrophagic markers. (Fig. 3B,D;Sup Fig. 3B, D)(Mittar et al., 2011) (Hogg et al., 1984), (van Lochem et al., 2004), (Abeles et al., 2012), (Hristodorov et al., 2015). Next, we observed that the autophagy inhibitors abrogated the functional parameters of activated macrophages (Fig. 3C,E; Sup Fig. 3C-E). Inflammatory cytokines viz. IL6, TNF-*a* and IL8 were assayed using semi-quantitative RT PCR and their production was severely downregulated in the presence of the inhibitors (Fig. 3D,Sup Fig. 3D).We further validated these results in differentiating PBMCs (Sup Fig. 4).Therefore, we could conclude that autophagy is extremely essential for promoting monocyte to macrophage differentiation.

**Fig. 3.**
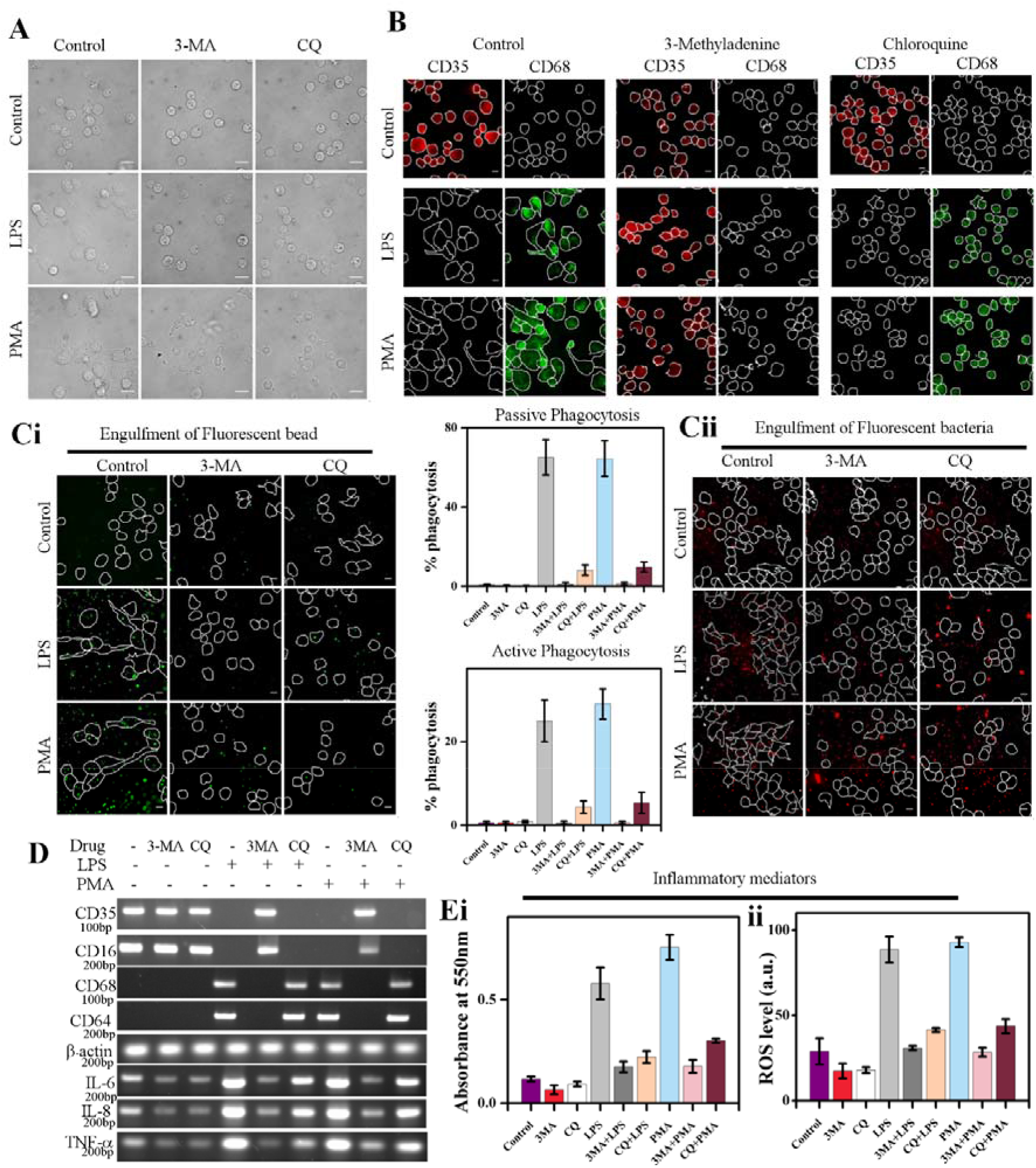
Autophagy is necessary to drive the process of monocyte differentiation. (A) DIC images of control (upper row), LPS (middle row) and PMA (lower row) stimulated THP-1 cells in presence of 3-Methyl Adenine (middle column) and Chloroquine (right column) 4 days post seeding. (B) Immunostaing of CD35 (left column) and CD 68 (right column) of THP-1 cells treated with 3-Methyl adenine and chloroquine in unstimulated conditions (upper row) and under stimulation/seeding with LPS (middle row), PMA (lower row). (C) Fluorescent images of passive (with fluorescent polystyrene bead) (i) and active phagocytosis (with RFP expressing *E.coli* (ii) of THP-1 cells under different conditions of stimulation in presence of 3-Methyl Adenine and Chloroquine. Graphical representation of the percentage of both passive and active phagocytosis is represented in the middle of (i) and (ii). (D) Semi-quantitative RT PCR data of monocyte/macrophage markers viz. CD35, CD16, CD68, CD64and inflammatory cytokines viz. IL-6, IL-8, TNF-α of THP-1 cells under different conditions of stimulation in presence of 3-Methyl Adenine and Chloroquine. Actin is used as control. (E) The quantitative comparison of NO (i) and ROS (ii) production fromTHP-1 cells under different conditions of stimulation in presence of 3-Methyl Adenine and Chloroquine. Scale Bar 10μm. Graphical analyses are performed with standard error bars after repeating the experiment at least thrice.

Next,to examine whether autophagy is also a crucial regulator of monocyte to macrophage differentiation *in vivo*, we injected 3-MA and CQ into the tail vein of mice followed by injection of RFP-*E.coli* into the peritoneal cavity of these mice.After 48 hours the mice were euthanized, the peritoneal fluid was extracted and total cell enumeration was performed. We found a sharp decline in the fluorescence intensity in the drug-treated mice, indicating a decreased recruitment of macrophages into the peritoneal cavity (Sup Fig. 5A). On spreading equal volumes of the peritoneal fluid on bacterial plates, we observed that peritoneal fluid extracted from drug-treated mice gave rise to a lawn of bacterial colonies (Sup Fig. 5B). Thus, we could effectively infer that a lesser number of macrophage infiltration due to significantly impaired autophagy resulted in the increment of bacterial titer on the culture plates. Moreover, a notable reduction in CD68 and CD64 expression in the drug-treated mice gave further credence to our claim (Sup Fig. 5C). A pronounced reduction in the expression of inflammatory cytokines and severely decreased levels of ROS and NO in the drug-treated micestrongly co-related with a sharp decline in macrophage recruitment(Sup Fig. 5D).Lastly, to determine whether the intensity of the immune regulation in response to bacterial infection was restricted to the peritoneum or it was a systemic phenomenon, we extracted the spleen from the animals and performed a comparative study of their sizes, indicative of the degree of infection, immune-cell infiltration and consequent inflammation. Spleen sizes of mice treated with drugs were found to be much smaller, further ratifying our hypothesis (Sup Fig. 5E).

### 3.4 Autophagy is not sufficient, but assists to drive the process of monocyte to macrophage differentiation

Next, we were curious to learn whether autophagy was sufficient to drive the process of monocyte differentiation.Monocytes were treated with certain inducers of autophagy, namely Rapamycin, which is known to induce autophagy via an mTOR-dependent mechanism and Sodium Valproate, which directs autophagy via an mTOR-independent pathway(Sarkar, 2013)in the absence of any chemical stimuli and subsequent morphology and marker analysis was performed. Suspended spherical morphology and the presence of monocytic markers confirmed that monocytes failed to differentiate (Fig. 4A and B). So,on observing that autophagy was not sufficient to drive monocyte differentiation, we examined whether it played an important role in assisting the process of differentiation. We found that simultaneous treatment of rapamycin along with a sub-optimal dose of PMA;reported to arrest cell cycle but not induce monocyte differentiation(Traore et al., 2005), resulted in successful monocyte differentiation (Fig. 4C-G). Therefore our data represent that, although autophagy cannot induce monocyte to macrophage differentiation it certainly plays a crucial role in promoting the process.

**Fig. 4.**
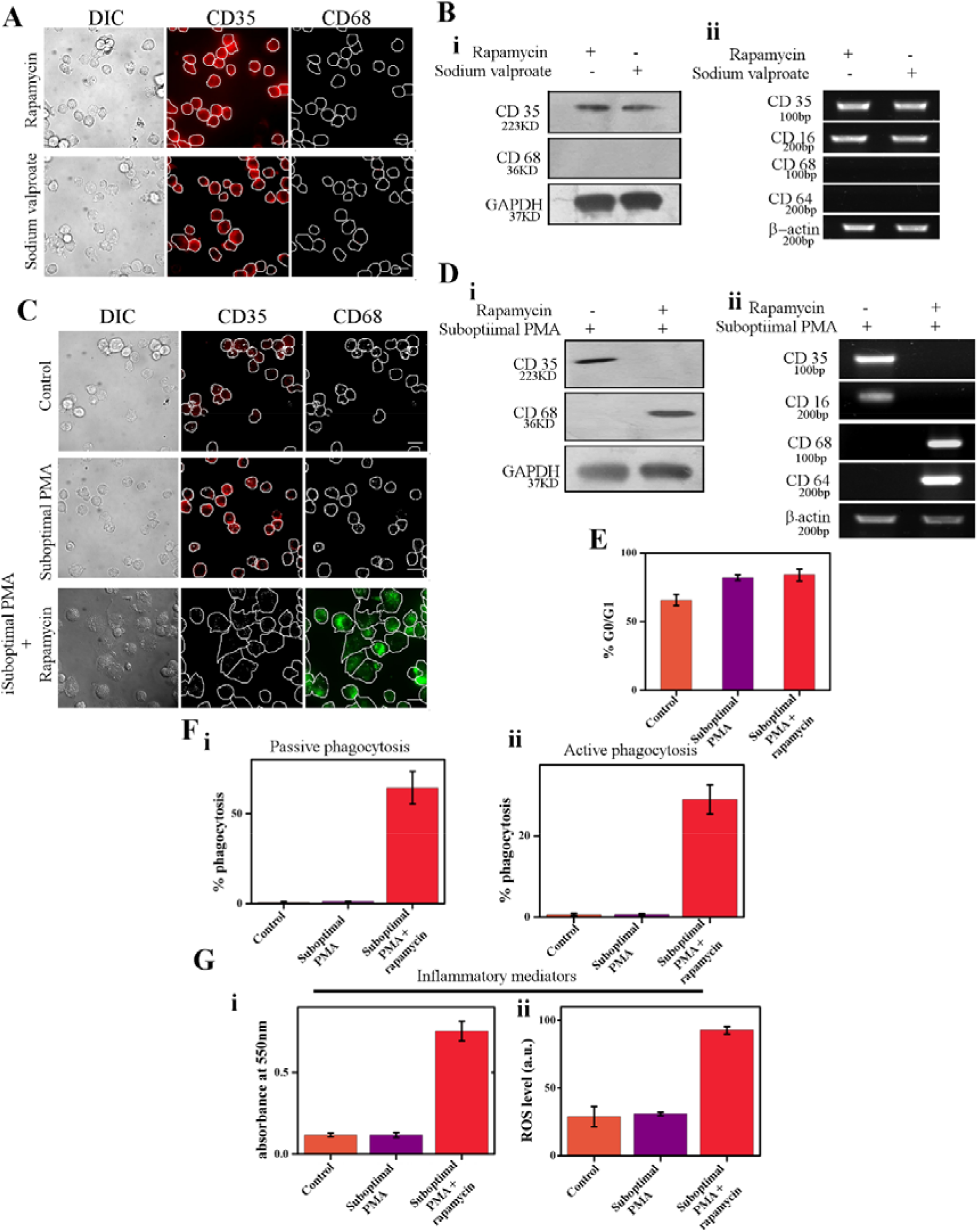
Autophagy is not sufficient, but assists to drive the process of monocyte to macrophage differentiation. (A) DIC (left panel) and immunofluorescence images of CD 35 (middle panel) and CD68 (right panel) of Rapamycin (upper) and Sodium Valproate (lower) treated THP-1 cells. (B) Western Blot of CD35 and CD68 (i) and semi-quantitative RT-PCR analysis of CD35, CD16, CD68 and CD 64 (ii) after treatment with Rapamycin/Sodium Valproate. GAPDH and Actin are used as controls respectively. (C) DIC (left panel) and immunofluorescence images of CD35 (middle panel) and CD68 (right panel) of control (upper), minimal PMA (middle) and Minimal PMA+ Rapamycin treated THP-1 cells. (D) Western Blot of CD35 and CD68 (i) and semi-quantitative RT-PCR analysis of CD35, CD16, CD68 and CD64 (ii) in THP-1 cells stimulated by a sub-optimal dose of PMA in the absence and presence of Rapamycin treatment. GAPDH and Actin are used as controls respectively. **(E)** Graphical representation of the percentage of cell-cycle arrest (G0/G1) in control, minimal PMA and minimal PMA + Rapamycin treated THP-1 cells. (F) Graphical representation of the percentage of passive (i) and active (ii) phagocytosis in control, minimal PMA and minimal PMA+ Rapamycin treated THP-1 cells. (G) The quantitative comparison of NO (i) and ROS (ii) production in control, minimal PMA and minimal PMA+ Rapamycin treated THP-1 cells. Scale Bar 20μm. Graphical analyses are performed with standard error bars after repeating the experiment at least thrice.

### 3.5 External ATP supplementation can overcome inhibited differentiation in monocytes with compromised autophagy

Next, we investigated the contribution of autophagy during monocyte differentiation. As autophagy is a major catabolic process of the cell, (Glick et al., 2010),(Singh and Cuervo, 2011),(Sakaki et al., 2013),(Diskin and Pålsson-McDermott, 2018),(Wang et al., 2018)we contemplated ATP (energy) generation to be one of the cardinal purposes of autophagy during monocyte differentiation. To verify our hypothesis, we quantified the total cellular ATP levels. We observed a steady increase in intracellular ATP levels in differentiating monocytes till Day 5 post-stimulation (Fig. 5Ai). Next, we assessed the intracellular ATP levels in chemically stimulated cells in the presence of autophagy inhibitors. The data suggested a significant reduction in ATP production in these inhibitor-treated cells (Fig. 5Aii, Sup Fig. 6A), implying that ATP generation is an important function of autophagy. In this particular scenario, we were curious to learn if supplementation of ATP from external sources could re-program the stalled differentiation process. Interestingly, we observed that treatment with external ATP could not induce spontaneous differentiation in monocytes (Fig. 5C-E). So much so, it was even unable to induce differentiation in 3-MA treated cells (Fig. 5B, 5F-K). But ATP supplementation could induce normal cellular spreading characteristic of macrophages, in CQ, Baf and Rab7 DN treated stimulated cells (Fig. 5B, Sup Fig. 6B).ATP treatment could also revertthe functional attributes of these cells (Fig. 5F-K, Sup Fig. 6C-E).

**Fig. 5.**
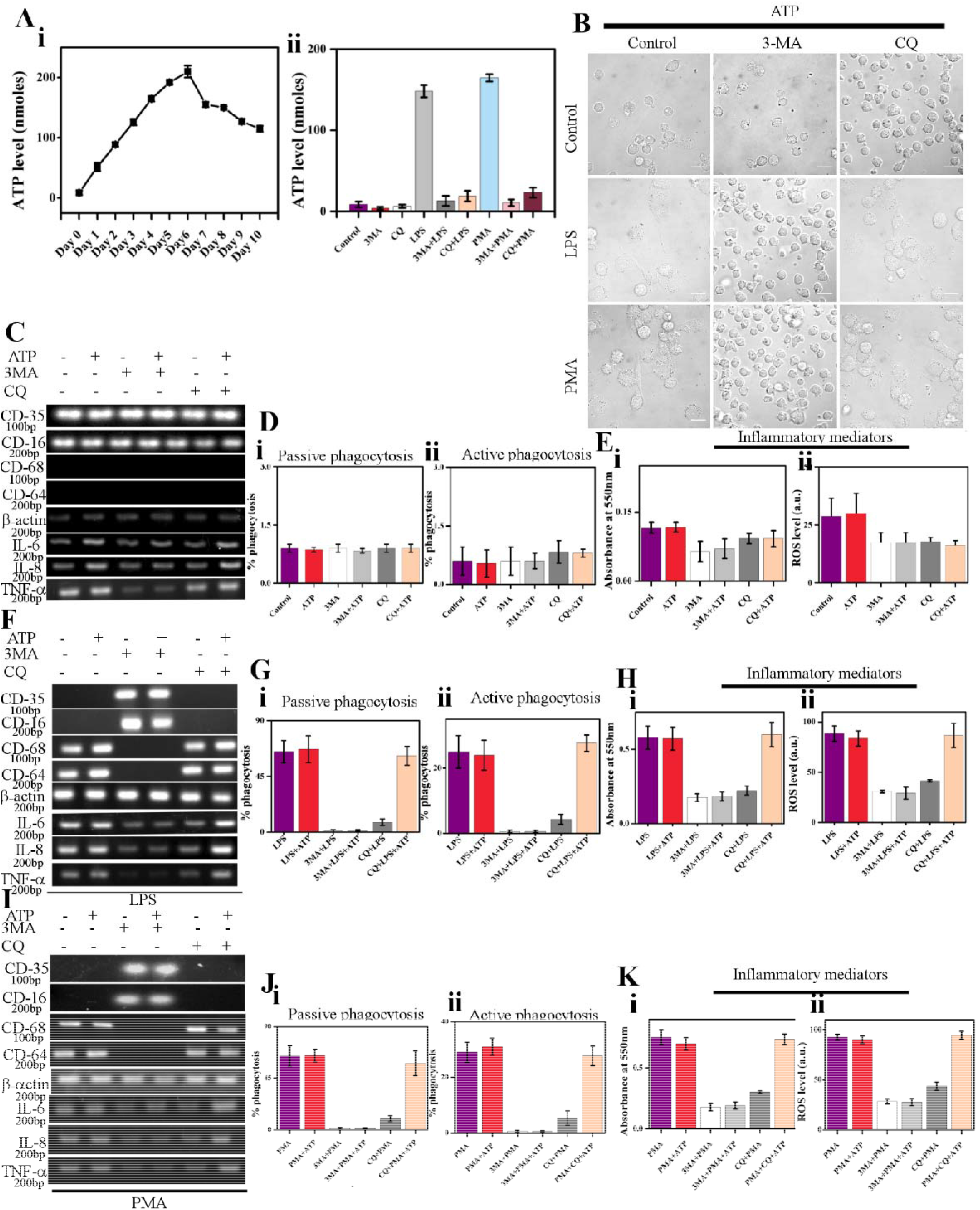
External ATP supplementation can overcome inhibited differentiation in monocytes with compromised autophagy. (A) Graphical representation of the quantification of total intracellular ATP levels vs. the no. of days (i) and under different conditions of stimulation in the presence of 3-Methyl Adenine and Chloroquine (ii) in differentiating THP-1 cells. (B) DIC images of untreated (left panel), 3-Methyl Adenine (middle panel) and Chloroquine (right panel) treated and external ATP supplemented control (upper), LPS (middle) and PMA (lower) stimulated THP-1 cells. (C) Semi-quantitative RT PCR analysis of monocyte/macrophage markers viz. CD35, CD16, CD68, CD64and inflammatory cytokines viz. IL-6, IL-8, TNF-α under varying conditions of 3-Methyl Adenine and Chloroquine treatment and ATP addition. Actin is used as a control. (D) Graphical representation of the percentage of passive (i) and active (ii) phagocytosis of THP-1 cells under different conditions of 3-Methyl Adenine and Chloroquine treatment and ATP addition. (E) The quantitative comparison of NO (i) and ROS (ii) production fromTHP-1 cells under different conditions of 3-Methyl Adenine and Chloroquine treatment and ATP addition. (F) Semi-quantitative RT PCR data of monocyte/macrophage markers viz. CD35, CD16, CD68, CD64and inflammatory cytokines viz. IL-6, IL-8, TNF-α under varying conditions of 3-Methyl Adenine and Chloroquine treatment and ATP addition along with LPS stimulation. Actin is used as a control. (G) Graphical representation of the percentage of passive (i) and active (ii) phagocytosis of THP-1 cells under different conditions of 3-Methyl Adenine, Chloroquine treatment and ATP addition along with LPS stimulation. (H) The quantitative comparison of NO (i) and ROS (ii) production fromTHP-1 cells under different conditions of 3-Methyl Adenine, Chloroquine treatment and ATP addition along with stimulation. (I) Semi-quantitative RT PCR data of monocyte/macrophage markers viz. CD35, CD16, CD68, CD64and inflammatory cytokines viz. IL-6, IL-8, TNF-α under varying conditions of 3-Methyl Adenine and Chloroquine treatment and ATP addition along with PMA stimulation. Actin is used as a control. (J) Graphical representation of the percentage of passive (i) and active (ii) phagocytosis of THP-1 cells under different conditions of 3-Methyl Adenine, Chloroquine treatment and ATP addition along with PMA stimulation. (K) The quantitative comparison of NO (i) and ROS (ii) production fromTHP-1 cells under different conditions of 3-Methyl Adenine, Chloroquine treatment and ATP addition along with PMA stimulation. Scale Bar 20μm. Graphical analyses are performed with standard error bars after repeating the experiment at least thrice.

Next, we investigated the underlying mechanism behind this ATP-induced reprogramming of morphological and functional parameters in CQ, Bafilomycin treated and Rab7 DN infected chemically stimulated cells. We found that supplementation of extracellular ATP fulfilled the intracellular dearth of ATP in these inhibitor-treated, chemically stimulated cells (Sup Fig. 7, 8). Moreover, we found that this enhanced level of intracellular ATP can overcome the inhibited fusion of the autophagosome with the lysosome (Sup Fig. 7, 8). Overall our data suggest that external ATP supplementation can fulfill the energy dearth of differentiating monocytes and at the same time is capable of inducing the stunted maturation of autophagy to sustain the process and hence the differentiation.

### 3.6 Autophagy dictates degradation of monocyte-specific proteins during differentiation

Many literature reports suggest that the protein profile of monocyte is significantly different from that of a macrophage(Liu. et al., 2008; Tuomisto et al., 2005).To analyze the qualitative protein content of a differentiating monocyte we estimated the total cell protein at different time points post-induction. Immediately after the addition of chemical stimuli, the total cell protein increased significantly till 1 hour and then there was a steady decline up to 8 hours followed by an effective increase. Therefore, the total cell protein of a differentiating monocyte follows a very unique profile that must be correlated with the cellular physiology. Thus, we assumed that protein degradation machineries could be playing a pivotal role in the differentiation process by regulating the cellular proteomic profile (Fig. 6A, Sup Fig. 9A).Ubiquitin-proteasomal pathway and autophagy are the major protein degradation pathways that help maintain a functional cellular proteome or proteostasis. We investigated the individual contributions of these two pathways during monocyte to macrophage differentiation. We treated monocytes with heclin, an E3 ubiquitin-ligase inhibitor before stimulation with LPS/PMA. Surprisingly, heclin failed to impart any significant change and the monocytes underwent successful differentiation as was confirmed by morphology and marker analysis (Fig. 6B-D). Finally, we quantified the total cell protein in chemically stimulated monocytes in the presence of various inhibitors. We found that heclin did not impart any significant difference on the overall protein degradation, indicating that the proteasomal pathway of protein degradation may not have a pronouncedimpact on monocyte differentiation.On the contrary; drugs like 3-MA and chloroquine had a profound impact on the protein profile of differentiating cells (Fig. 6E). So in thenext step, we analyzed whether autophagy serves as a crucial protein degrading machinery to achieve the differential protein profile that characteristically distinguishes macrophages from monocytes.We have already established (Fig. 3), that pharmacological inhibitors of autophagy have a notable impact on monocyte differentiation. To investigate the contribution of autophagy in cellular protein degradation in a differentiating monocyte, we used chloroquine, bafilomycin and Rab7DN adenoviral construct, which are known to block the maturation step of autophagy. In response to LPS/PMA treatment, CD35 was found to retain till 6 hours post-treatment. However, in the presence of these inhibitors CD 35 was found to accumulate in the cytosol in puncta form (Fig. 6F, Sup Fig. 9D). Surprisingly some of these puncta were found to co-localize with LC3-II puncta, implying that autophagy serves as the protein degradation machinery in differentiating monocytes (Fig. 6F, Sup Fig. 9D).Next, to determine whether autophagy regulates the amino acid recycling in differentiating monocytes,we labeled monocyte cellsurface proteins with cell-impermeable NHS Biotin before stimulation. After differentiation, cells were lysed and biotinylated proteins were pulled down using streptavidin beads. Next, pulled down sample was probed with anti CD 68 antibody in Western Blot. CD 68 is a cellsurface marker that is exclusively present in macrophages and was selected for its ‘all-but-none’ type expression in monocyte vs macrophagic condition. The presence of CD68 in the pulled-down fraction indicated that biotinylated amino acids present in monocyte proteins were reused for the synthesis of new proteins in the differentiating cells (Fig. 6G). To validate the role of autophagy in the substrate recycling pathway, we did similar experiments in the presence of autophagy inhibitors. Incidentally, we could not detect the CD 68 band in the pulled-down sample (Sup Fig. 9C and E) and therefore, this underscores the claim that autophagy is the master regulator of the protein degradation and substrate recycling that ultimately guides monocytedifferentiation. Thus ATP generation and substrate recycling were identified as the two important functions of autophagy during monocyte to macrophage differentiation. Next, we aimed to determine whether autophagy could support the differentiation process independently i.e. in absence of external nutrients. So, stimulated monocytes were seeded in minimal buffer media like EBSS and HBSS. We observed the effective differentiation of monocytes under starved conditions (Fig. 6H, Sup Fig. 9F). Further, to check whether substrate recycling driven monocyte differentiation in starved condition proceedsvia the protein degradation pathway, we used various inhibitors. Heclin imparted no observable change but differentiation in starved condition was found to be blocked by CQ, Bafilomycin and Rab7 DN (Fig. 6H, Sup Fig. 9F-H). Thus overall our data suggest that autophagy functions as a major substrate recycling machinery to drive the process of monocyte differentiation.

**Fig. 6.**
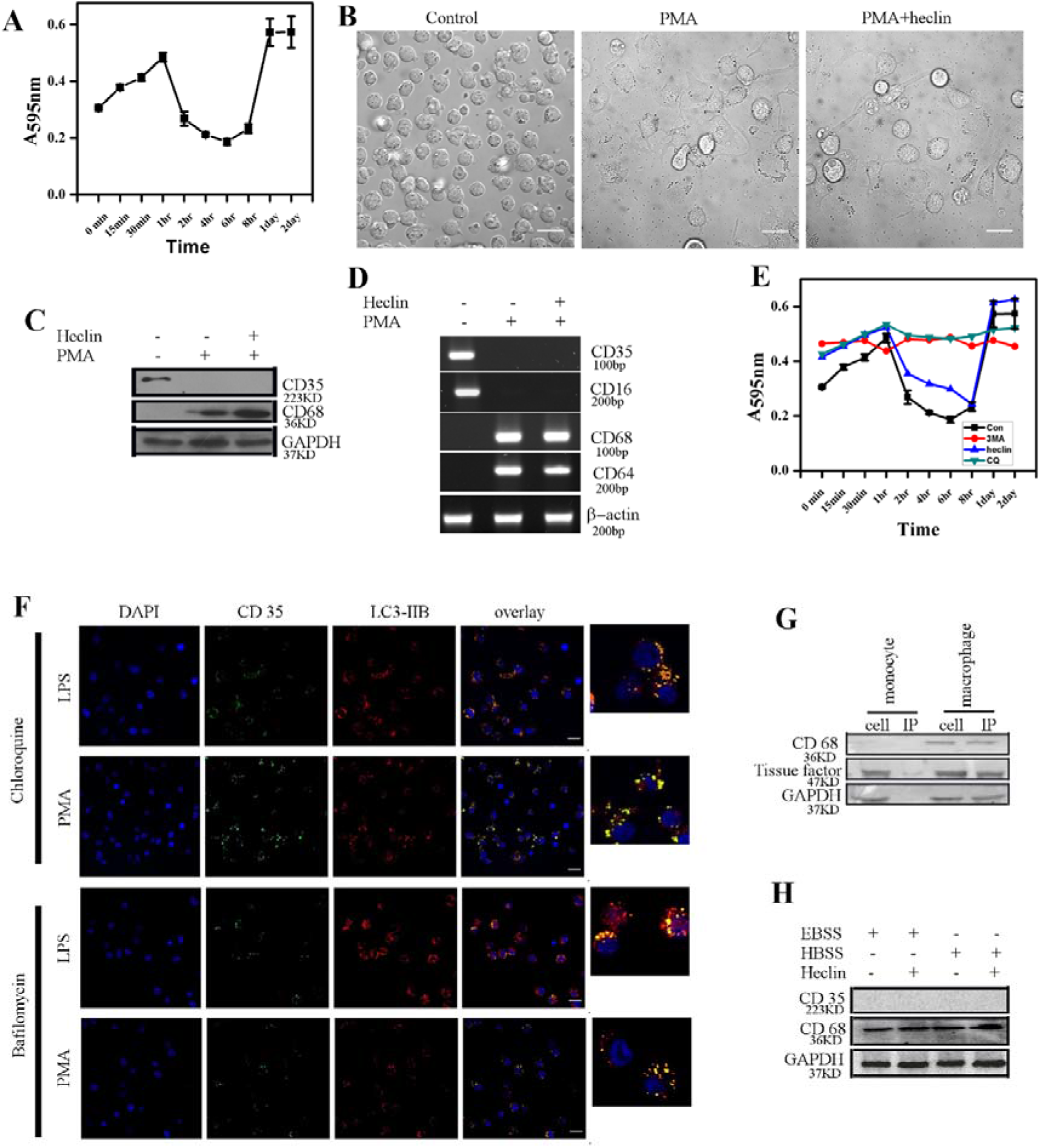
Autophagy dictates degradation of monocyte-specific proteins during differentiation. (A) Graphical representation of the quantitative estimation of total cell protein of THP-1 cells at different time points post stimulation with PMA. (B) DIC images of THP-1 cells in unstimulated (left) and PMA stimulated conditions in absence (middle) and presence of Heclin (right). (C) Western Blot of CD35 and CD68 in THP-1 cells in unstimulated and PMA stimulated conditions in presence and absence of Heclin. GAPDH is used as a loading control. (D) Semi-quantitative RT PCR analysis of monocyte/macrophage markers viz. CD35, CD16, CD68, CD64 in unstimulated and PMA stimulated conditions in presence and absence of Heclin. Actin is used as a control. (E) Graphical representation of quantitative estimation of total cell protein of THP-1 cells at different time point post stimulation with PMA in presence of 3-Methyl adenine (3MA), Chloroquine (CQ) and Heclin. (F) DAPI stained (1^st^column)and immuno-fluorescence images of CD35 (2^nd^column) and LC3-IIB (3^rd^column) stained LPS (upper) and PMA (lower) stimulated Chloroquine (upper two panels) and Bafilomycin (lower two panels) treated THP-1 cells. The 4^th^column represents the superimposed (overlay) images of the 2^nd^ and 3^rd^ panels. Magnified and superimposed images are presented at the extreme right of the panel. (G) Western Blot of CD68 following affinity pull-down using streptavidin bead in monocyte and macrophage fractions. Tissue Factor and GAPDH are used as loading controls. (H) Western blot of CD35 and CD68 after seeding THP-1 cells in EBSS & HBSS minimal media in the presence or absence of Heclin. GAPDH is used as the loading control. Scale Bar 25 μm.

### 3.7 Autophagy regulates cell-cycle during monocyte to macrophage differentiation

Autophagy is an intracellular degradative process and has been widely reported to downregulatetumorigenesis by suppressing the proliferation of cancer cells(Byun et al., 2017; Jing et al., 2014),(Kondapuram et al., 2019). We were curious to understand whether a similar mechanism operates in the case of differentiating monocytes and investigated the complex cross-talk between autophagy, cell-cycle arrest and differentiation. Monocytes were grown in the presence of Rapamycin and Phosphatidic acid and the total cell count was documented 48 hours post-treatment. Interestingly, we observed that the total cell-count increased significantly in the Phosphatidic acid treated population. However, there was a sharp depreciation in the total cell-count in the Rapamycin treated set (Fig. 7A) although there was no cell-cycle arrest. Moreover, it is reported that inducer treatment leads to the arrest of the monocyte cell cycle before differentiation (Traore et al., 2005).Our previous study had also confirmed that LPS and PMA arrest the cell cycle of THP-1 cells at G0/G1 step during monocyte differentiation(Bhattacharya et al., 2018).Thus to assay the impact of autophagy in cell cycle arrest of a differentiating macrophage, we analyzed cell cycle and cellular proliferation in the presence of different autophagy inhibitors. Surprisingly, no cell cycle arrest was detected and normal proliferation of monocytes was observed when the monocytes were treated with PA or 3MA (Known to block initial stages of autophagy) before the stimulation with LPS/PMA (Fig. 7B). Therefore, we concluded that inhibition of autophagy failed to arrest the cell-cycle during monocyte to macrophage differentiation. Thus, this highlights the fact that autophagy is required to bring about cell-cycle arrest in monocytes. To further our understanding, we carried out proteomic profiling of monocytes, autophagosomes and macrophages using MALDI-MS and LC-MS/MS. Consistent with our observations so far, we identified a list of proteins that are involved in cell-cycle progression in the autophagosomes as well as in monocytes. (Fig. 7C)Cell-cycle arrest prior to differentiation is well reported. Consistent with this notion, our data also reflects the fact that the proteins involved in cell-cycle progression get degraded by autophagy during the course of monocyte differentiation.

**Fig. 7.**
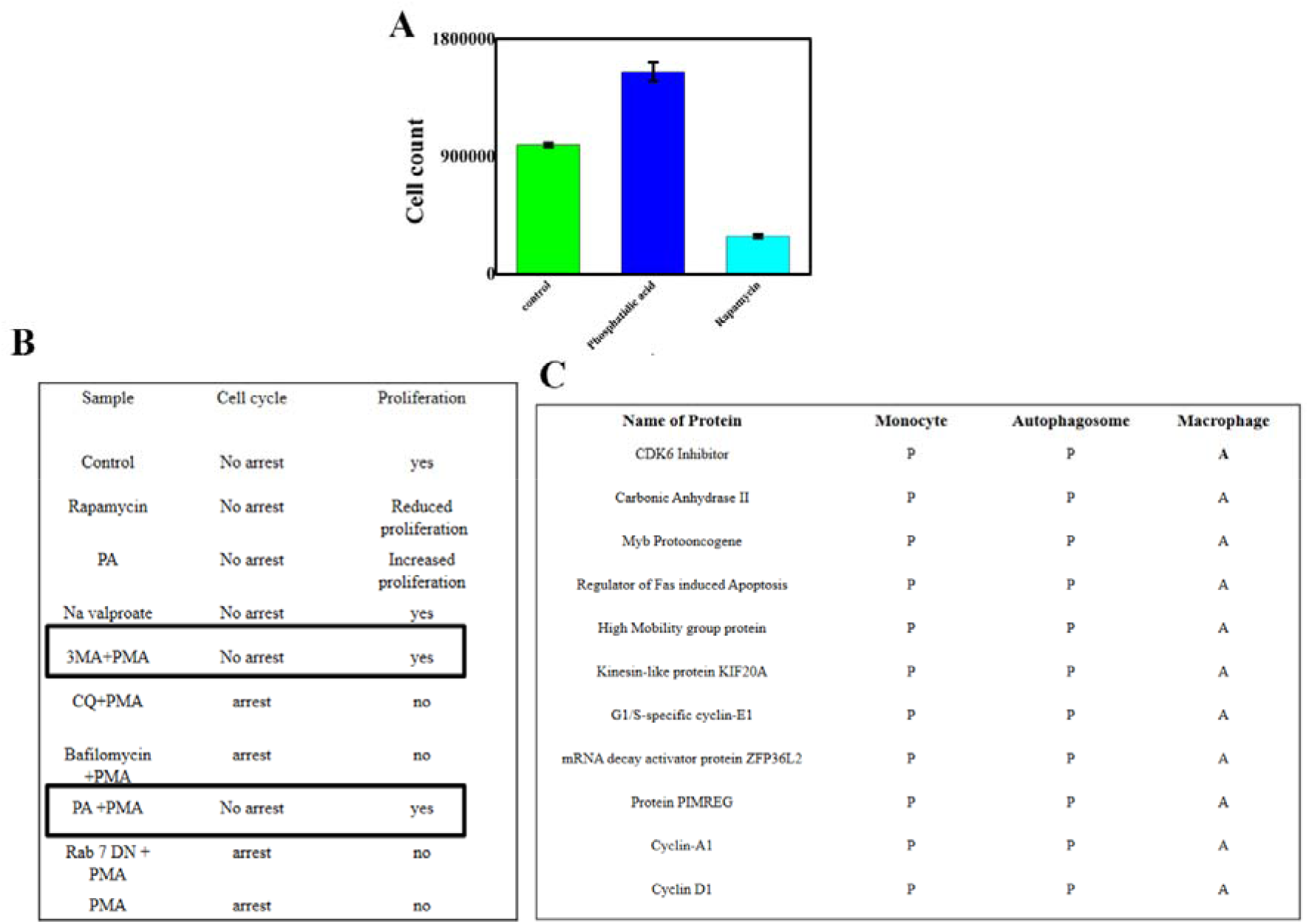
Autophagy regulates cell-cycle during monocyte to macrophage differentiation. (A) Graphical estimation of total cell count 48 hours post-treatment in control, phosphatidic acid and rapamycin treated THP-1 cells. (B) Tabular documentation of cell cycle arrest and extent of cellular proliferation in THP-1 cells under different conditions of treatment and stimulation. (C) Tabular documentation of the presence/absence of selected proteins in the monocyte, autophagosome and macrophage fractions.

### 3.8 Adhesion is essential for autophagy maturation during monocyte differentiation

We had previously established that adhesion is indispensable for monocyte differentiation in 2D/3D micro-environment(Bhattacharya et al., 2018). As adhesion is central to monocyte differentiation, we were intrigued to learn whether it was vital for autophagy.For this, chemically stimulated monocytes were seeded on an adhesion-incompatible agarose bed. Although our data confirm the induction of autophagy in the absence of adhesion, maturation was abrogated on the adhesion-incompatible surface (Fig. 8A and B). Next, the cells were recovered from the agarose bed, washed and seeded on an adhesion-compliant surface; resulting in autophagy maturation and completion of the differentiation process (Fig. 8D and E). Previous work from our group had indicated the importance of NFκβ-mediated adhesion on monocyte differentiation(Bhattacharya et al., 2018). Since we observed that adhesion is a crucial parameter for regulating autophagy maturation, we were curious to examine the role of NFκβ in monitoring the fusion of lysosome with the autophagosome. Cells were similarly recovered from the agarose bed and incubated in the presence of NFκβ Inhibitor (NFκβI) on an adhesion-compatible surface. Autophagy maturation was found to be halted in NFκβI treated monocytes and the cells failed to differentiate and stained positive for monocyte-specific markers (Fig. 8F-G).Taken together, these data highlight the fact that NFκβ mediated cellular adhesion is essential for autophagy maturation.

**Fig. 8.**
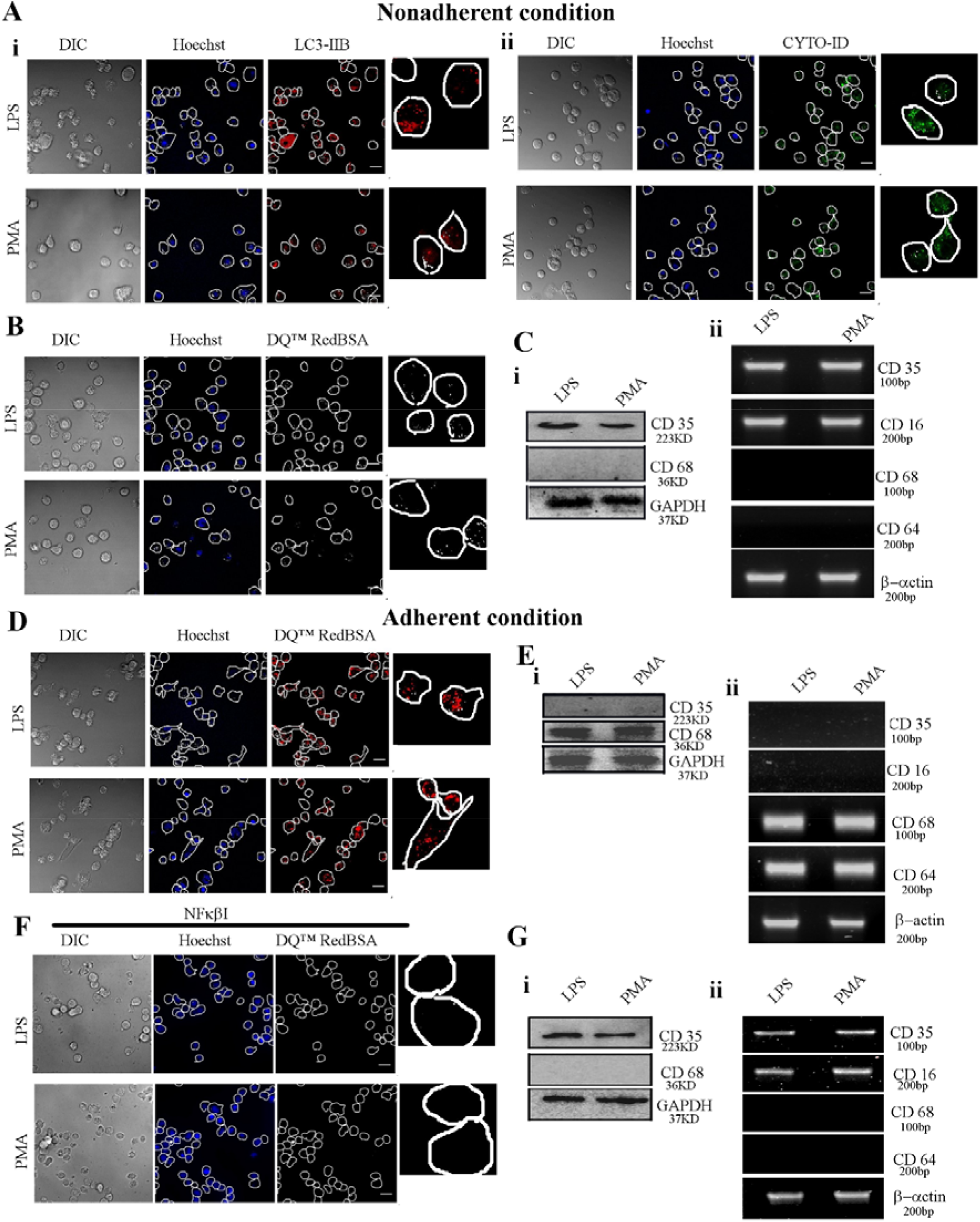
Adhesion is essential for autophagy maturation during monocyte differentiation. (A) (i) DIC (left panel) and fluorescent images of Hoechst stained (middle panel) and LC3-IIB (right panel) immunostained LPS (upper) and PMA (lower) stimulated THP-1cells under non-adherent condition. Magnified view of LC3-IIBimmunostained cells are represented at the right side of the panel. (ii) DIC (left panel) and fluorescent images of Hoechst stained (middle panel) and CYTO-ID (right panel) stained LPS (upper) and PMA (lower) stimulated THP-1 cells under non-adherent condition. Magnified view of CYTO-ID stained cells are represented at the right side of the panel. (B) DIC (left panel) and fluorescent images of Hoechst (middle panel) and DQ™ Red BSA (right panel) stained LPS (upper) and PMA (lower) stimulated THP-1cells under non-adherent conditions. Magnified view of DQ™ Red BSA stained cells are represented at the right side of the panel. (C) Western Blot of CD35 and CD68 (i) and semi-quantitative RT-PCR analysis of monocyte/macrophage markers CD35, CD16, CD68 and CD64 (ii) of THP-1 cells under non-adherent conditions. GAPDH and Actin are used as controls respectively. (D) DIC (left panel) and fluorescent images of Hoechst (middle panel) and DQ™ Red BSA (right panel) stained LPS (upper) and PMA (lower) stimulated THP-1cells under adherent conditions. Magnified view of DQ™ Red BSA stained cells are represented at the right side of the panel. (E) Western Blot of CD35 and CD68 (i) and semi-quantitative RT-PCR analysis of monocyte/macrophage markers CD35, CD16, CD68 and CD64 (ii) of THP-1 cells under adherent conditions. GAPDH and Actin are used as controls respectively. (F) DIC (left panel) and fluorescent images of Hoechst stained (middle panel) and DQ™ Red BSA (right panel) immunostained LPS (upper) and PMA (lower) stimulated THP-1cells in presence of NFκβ Inhibitor (NFκβI). (G) Western Blot of CD 35 and CD 68 (i) and Semi-quantitative RT-PCR analysis of monocyte/macrophage markers CD35, CD16, CD68 and CD64 (ii) in the presence of NFκβI. GAPDH and Actin are used as controls respectively.

## 4. Discussion

The mononuclear phagocytic system comprising of monocytes and macrophages primarily links innate and adaptive immune systems. Although theimportance of monocyte to macrophage differentiation is highly celebrated in physiology, the underlying mechanistic details remain elusive.In this manuscript, we have correlated autophagy with the process of monocyte to macrophage differentiation andelucidated the detailed role of autophagy in this context. Here, we have also demonstrated the active participation of mTOR dependent classical autophagy pathway during monocyte to macrophage differentiation.

Autophagy i.e. the process of self-eating was previously considered detrimental leading to cellular death (Bialik et al., 2018). But growing evidence has rejected the antiquated concept of autophagic cell death and in the present scenario, it is considered as a crucial regulator of cellular homeostasis (Raquel et al., 2013),(Chun and Kim, 2018; Ryter et al., 2013),(G. Kroemer, 2008). In recent past, autophagy has been correlated with several cellular physiological processes including diseased conditions like cancer, cell death, neurodegenerative diseases, Type II diabetes, fatty liver, infectious diseases, cardiomyopathies, cellular aging and innate immune system(R. Mathew, V.K-Wadsworth, 2007),(Yonekawa and Thorburn, 2013),(Nixon, 2013),(Yang et al., 2017),(Czaja, 2016),(Desai et al., 2015),(Barbosa et al., 2019). In this manuscript, we have correlated autophagy with the process of monocyte to macrophage differentiation andelucidated the detailed role of autophagy in this context.

Primarily, the results of this study show that autophagy is induced ubiquitouslyduring monocytedifferentiation irrespective of the mode of induction. This observation is also supported by many otherindependent reports(Jacquel et al., 2012; Yuk et al., 2009),(Zhang et al., 2012); however, those reports do not adequately depict the intricate detailing necessary to successfully establish the complex relationship between autophagy and monocyte differentiation. Although various chemical stimuli like GMCSF, CSF, IL27, LPS, PMA are reported to induce autophagy during monocyte differentiation, the molecular mechanism behind autophagy was not explainedpreviously. Here, we demonstrate the active participation of mTOR dependent classical autophagy pathway during monocyte to macrophage differentiation.

Zhang et al and Jacquel et al have shown autophagy to be intrinsically associated with the monocyte differentiation process(Zhang et al., 2012)(Jacquel et al., 2012; Jacquel et al., 2016). To further analyze the contribution of autophagy in the process of monocyte differentiation, we investigated the consequence of inhibiting autophagy by pharmacologic and other genetic manipulation approaches. 3-methyladenine (3-MA), a Vps34 inhibitor was found to abrogate the monocyte differentiation procedure. On the contrary, treatment with Chloroquine, Bafilomycin and infection with the adenoviral construct of Rab 7 DN partially affected the differentiation process. Although these cells adhered to the surface and expressed macrophage-specific markers, they retained the spherical morphology. Moreover, their phagocytic and inflammatory potentials were seriously compromised.Importantly, 3-MA abolished LC3-II processing and CQ, Baf, Rab7 DN affected the autophagosome maturation process. We performed similar experiments using an animal model (mice) and the results further endorsed our proposition and therefore we successfully established thatautophagy is essential for the production of functional macrophages.

In this study, we demonstrate that autophagy is not sufficient to induce monocyte differentiation on its own but can definitely assist the process. There are several autophagy-independent processes that guide monocyte differentiation. Martinez et al have performed an in-depth transcriptional profiling of genes that are differentially regulated during monocyte differentiation (Martinez et al., 2006). Their data identifies differential regulation of genes involved in immune response, DNA metabolism, cell cycle, membrane receptors, lipid metabolism, chemotaxis and G protein coupled receptors, macromolecule biosynthesis, protein metabolism, carbohydrate metabolism, phosphate metabolism as critical factors for monocyte differentiation. Differentiation is also accompanied by the accretion in the number of cytosolic organelles like lysosome, mitochondria, etc essential for acquirement of augmented phagocytic ability by differentiating macrophages (Cohn and Belinda, 1965) (Daigneault et al., 2010) (Cohn et al., 1966). In our previous work, we had established actomyosin dynamics as a crucial parameter for sustaining monocyte differentiation (Bhattacharya et al., 2020). Traore et al had successfully used a suboptimal dose of PMA which was unable to sustain the differentiation process but could successfully induce cell cycle arrest(Traore et al., 2005). Here, we have shown that co-administration of Rapamycin along with that specific suboptimal dose of PMA can complete the differentiation process. Thus we concluded that monocyte differentiation is a highly complex multi-faceted phenomena and autophagy is one of the crucial mediators of the process.

Studies investigating the molecular mechanisms of cross-talk between differentiation and autophagy are still in their infancy. There are very few reports that explain the contribution of autophagy in the context of differentiation. Zhang et al have shown that autophagy is a prerequisite for monocyte differentiation to delay apoptotic cell death(Zhang et al., 2012). However, in this manuscript, we have done an in-depth investigation of different functions of autophagy in the differentiation process. Some independent reports explain the crucial role of autophagy in maintaining cellular energetic balance(Lum et al., 2005; Singh. Rajat, 2012),(Glick et al., 2010),(Singh and Cuervo, 2011).Moreover, monocyte to macrophage differentiation is considered to be an energy-demanding process(Sakaki et al., 2013),(Diskin and Pålsson-McDermott, 2018),(Wang et al., 2018). Differentiation is accompanied by cellular spreading, cytoskeletal dynamics, synthesis of unique stage-specific biomolecules, enhanced production and secretion of cytokines, chemokines, interleukins, the enhanced genesis of cytoplasmic organelles, etc.Through this study, we identified autophagy as the energy-yielding machinery, supplying energy to satiate the energy-demand and fulfil the severe dearth of energy in monocytes during differentiation. Thus, supplementation with external ATP could successfully restore the stunted differentiation of monocytes treated with inhibitors that block the autophagy maturation process.

Autophagy is considered as a crucial regulator of nutrient recycling and the organelle repair process(He et al., 2018). Several groups have reported that autophagy primarily degrades the enclosed biomolecules like protein, lipid and carbohydrate and the digested simpler molecules (amino acid, simple sugar and lipids) are reused for further synthesis(Jun and Yoshinori, 2005),(Kenific et al., 2016),(Bhattacharya et al., 2018).Few reports suggest that the protein profile of macrophages is significantly different from that of monocytes (Dinnes et al., 2006; Portevin et al., 2015). We have found that the monocyte marker CD 35 gets degraded via autophagy. Several groups have reported a crucial regulatory role of autophagy whenever there is an excessive burden of proteins that need to be degraded (Ebrahimi-fakhari et al., 2011). Since differentiation is a robust process involving a massive turnover of proteins, we propose autophagy as the major protein degrading machinery during monocyte differentiation on the basis of our data. We have also shown autophagy-mediated recycling of the amino acids in differentiating monocytes. Finally, the successful differentiation of monocytes in the absence of any external supply of nutrients further confirms the crucial supportive role of autophagy in differentiation. Thus our study has unraveled a unique mechanistic correlation between autophagy and differentiation.

It is well reported that cell cycle arrest precedes monocyte to macrophage differentiation(Traore et al., 2005),(Rots et al., 1999). Traore et al have shown that PMA treatment leads to decreased phosphorylation of retinoblastoma protein with the concomitant downregulation of cdk4 (Traore et al., 2005). PMA treatment also results in PKC-ERK pathway-dependent up-regulation of P21(Traore et al., 2005). On the contrary, autophagy also has an inverse relationship with cellular proliferation(Azzopardi et al., 2016; Neufeld, 2012). Thus, Rapamycin and other synthetic autophagy inducers are clinically used as antiproliferative agents in the treatment of cancer(Byun et al., 2017; Jing et al., 2014; Kondapuram et al., 2019). Here, we have correlated autophagy with cell cycle arrest which is a prerequisite for monocyte differentiation. According to our present findings, autophagy specifically degrades the proteins responsible for cell-cycle progression to arrest the cell cycle resulting in the subsequent differentiation of monocytes to macrophages.

Kenific et al have established a detailed contribution of autophagy in cellular adhesion and migration(Kenific et al., 2016). Lee et al have also reported that loss of function of ATGs (autophagy-related gene) leads to defects in cellular adhesion-mediated cardiac morphogenesis in both zebrafish and mice embryos(Kenific et al., 2016). In this manuscript, we have also found that inhibition of autophagy affects monocyte adhesion as well as their differentiation to macrophages. Moreover, here we have explored the correlation between adhesion and autophagy from a reciprocal point of view. Previous publications from our group had shown that adhesion is necessary for monocyte differentiation(Bhattacharya et al., 2018). To date, there are no significant reports that analyze the effect of adhesion on autophagy. In this manuscript, we have established cellular adhesion to be a crucial regulator of autophagy. Moreover, we have explored the role of NFκβ in the context of autophagy maturation. Trocoli. Et al have reviewed the complex interrelation between autophagy and NFκβ in cancer cell (Ebrahimi-fakhari et al., 2011). In this manuscript, we furthered this understanding and established a strong correlation between cellular adhesion, NFκβ signaling and autophagy in the context of monocyte to macrophage differentiation.

Chloroquine, a potent inhibitor of autophagy, is a strong candidate for the treatment of rheumatoid arthritis (Jang et al., 2006). Although this treatment has long been in use, the underlying mechanistic details that contribute to the efficacy of this drug remain unknown and are probably believed to work by preventing the release of TNF and other pro-inflammatory cytokines from monocytes and macrophages (Jang et al., 2006). It is well-documented that macrophages produce larger amounts of cytokines as compared to monocytes(Scull et al., 2010). Our study provides strong evidence in favor of this particular observation. Administration of chloroquine is found to be beneficial in the treatment of rheumatoid arthritis, possibly because of its ability to inhibit the process of monocyte to macrophage differentiation by blocking autophagy.

Through this study, we have successfully established that autophagy is a dire necessity and an unavoidable need during monocyte to macrophage differentiation. In this manuscript, we have delved deeper into understanding the mechanistic complexities that interconnect autophagy with monocyte differentiation and our findings strongly indicate a crucial role of autophagy in this process. Overall, our study would provide a strong foundation for future studies on the role of autophagy in infectious, auto-immune and inflammatory diseases.

## Authorship Contribution

AB, DKS and PS designed the experiments; AB, PG, AS, AG and AB# (Arghya Bhowmick) performed the experiments and analyzed the data; AB, PG and PS wrote the manuscript. AG#(Abhrajyoti Ghosh)has helped in the experiment.

## Acknowledgment

PS acknowledges Grant No-SR/SO/BB-0125/2012 (Department of Science and Technology, India). We acknowledge CSIR for providing fellowship to AB (SPM). We acknowledge Debapriya Ghatak for helping us in using a confocal microscope and Tanojit Sur for helping us in FACS. We are indebted to Dr. Mahadeb Pal, Bose Institute, Kolkata for allowing us to use the Confocal Microscope facility.

